# Brain-engrafted macrophages provide protection against therapeutic irradiation and secondary concussive injury

**DOI:** 10.1101/794354

**Authors:** Xi Feng, Elma S. Frias, Maria S. Paladini, David Chen, Zoe Boosalis, McKenna Becker, Sonali Gupta, Sharon Liu, Nalin Gupta, Susanna Rosi

## Abstract

Brain resident microglia have a distinct origin compared to macrophages in other organs. Under physiological conditions, microglia are maintained by self-renewal from the local pool, independent of hematopoietic progenitors. Pharmacological depletion of microglia during therapeutic whole-brain irradiation prevents synaptic loss and long-term recognition memory deficits but the mechanisms behind these protective effects are unknown. Here we demonstrate that after a combination of therapeutic whole-brain irradiation and microglia depletion, macrophages originating from circulating monocytes engraft into the brain and replace the microglia pool. Comparisons of transcriptomes reveal that brain-engrafted macrophages have an intermediate phenotype that resembles both monocytes and embryonic microglia. Brain-engrafted macrophages display reduced phagocytic activity for synaptic compartments compared to microglia from normal brains in response to a secondary concussive brain injury. In addition to sparing mice from brain radiotherapy-induced long-term cognitive deficits, replacement of microglia by brain-engrafted macrophages can prevent concussive injury-induced memory loss. These results demonstrate the long-term functional role of brain-engrafted macrophages as a possible therapeutic tool against radiation-induced cognitive deficits.

## Introduction

Brain resident microglia are the innate immune cells of the central nervous system (CNS). Arisen from the yolk sac during embryonic development, microglia actively survey the environment to maintain normal brain functions (1, 2). Under physiological conditions, microglia are maintained solely by self-renewal from the local pool (3). Following brain injury and other pathological conditions, microglia become activated and play a central role in the clearance of cellular debris, but if not controlled this process can result in aberrant synaptic engulfment (4–7). Temporary depletion of microglia can be achieved by using pharmacologic inhibitors of the colony-stimulating factor 1 receptor (CSF-1R) (8). In the normal brain, treatment with CSF-1R inhibitors (CSF-1Ri) can deplete up to 99% of microglia without causing detectable changes to cognitive functions (8, 9). Full repopulation occurs within 14 days of inhibitor withdrawal and the repopulated microglia are morphologically and functionally identical to the microglia in naïve brains (9). Microglia depletion and repopulation by local progenitors has been shown to be beneficial for disease- ,injury- , and age-associated neuropathological and behavioral conditions(10–15). However, the mechanisms for these protective effects are unknown.

Whole-brain radiotherapy (WBRT), delivered in multiple fractions, is routinely used to treat patients with brain tumors. It is estimated that more than 200,000 patients receive WBRT yearly in the US alone (16). While it is effective in improving intracranial tumor control, WBRT leads to deterioration of patients’ cognitive functions and quality of life (17–19). Currently, there is no treatment available to prevent or mitigate these adverse effects. Previous studies demonstrated that WBRT causes deleterious effects to the CNS microenvironment by a number of mechanisms including apoptosis of neural progenitor cells, disruption of the blood-brain barrier, activation of microglia and accumulation of peripherally derived macrophages (20–25). We and others have reported that depletion of microglia during or shortly after brain irradiation in animal models can prevent loss of dendritic spines in hippocampal neurons and cognitive impairments that develop at later time points (12–14). These reports suggest that microglial plays a critical role in inducing synaptic abnormalities and consequently, cognitive deficits after brain irradiation. The underlying molecular pathways responsible for the protective effects of repopulated microglia against radiotherapy-induced neuronal alterations remain unknown.

In the current study, 1) we first defined signature of repopulating cells and analyzed the transcriptional profile of repopulated brain macrophages from irradiated mouse brains after CSF-1R inhibitor-mediated depletion. 2) We next sought to establish the origin of repopulated cells coming from the peripheral monocytic lineage. 3) We identified the functionality of repopulated macrophages by measuring the ability to engulf synaptic compartments compared to brain resident microglia. Lastly, 4) we determined the protective properties of brain-engrafted macrophages (BEMs) against a secondary concussive brain injury-induced cognitive deficits. Together, these results uncover the mechanism by which brain-engrafted macrophages preserve hippocampal synapses and memory functions after radiation injury and also in response to an additional brain injury.

## Significance Statement

This study reports fate and functions of brain-engrafted macrophages after they replaced resident microglia in the brain. Concurrent microglia depletion and therapeutic brain irradiation results in replacement of microglia by peripheral derived brain-engrafted macrophages, which maintain a stable population in the brain for at least six months. These brain-engrafted macrophages are not reactivated as irradiated microglia and do not exhibit irradiation-induced transcriptomic signatures. In addition, they express lower phagocytic and lysosome markers, and do not respond to a secondary concussive brain injury. As a result, long-term memory functions are protected in brain-engrafted macrophages bearing animals. We conclude that replacement of microglia by brain-engrafted macrophages protects against radiation- and concussive injury-induced memory deficits.

## Results

### Microglia depletion and repopulation prevents radiation-induced hippocampal-dependent memory deficits

Temporary microglia depletion during or shortly after exposure to brain irradiation prevents cognitive deficits, suggesting microglia’s key role in modifying neuronal and cognitive functions (12–14). Changes in expression levels of pro-inflammatory cytokine/chemokines have been shown to correlate with cognitive performance in mice (12, 15, 20), however, the exact change in the transcriptional profile of repopulated microglia after brain irradiation is unknown and is an important tool to dissect the roles that repopulated microglia play in preventing of radiation-induced memory deficits. In this study we used a previously reported WBRT and microglia depletion paradigm(13) and performed RNA sequencing using repopulated microglia with and without WBRT, and compared with transcriptomes of microglia obtained from mice without CSF-1R inhibitor-mediated depletion (Figure 1a,b). A CSF-1R inhibitor was used to fully deplete microglia in 8-weeks old male mice, for a duration of 21 days. Three fractions of therapeutic whole-brain irradiation were given to each mouse every other day over five days starting from day 7 of CSF-1R inhibitor treatment. Novel Object Recognition (NOR) test was used to measure recognition memory 4 weeks after the last fraction of WBRT. Consistent with our previous report, fractionated WBRT resulted in impairment in recognition memory, which was prevented by CSF-1R inhibitor treatment (Figure 1a**, lower panel**). No deficits in motor functions or changes of anxiety levels were found in the open field test (scored from day 1 of NOR test, data not shown). One day after the NOR test, mice were euthanized and whole brains were used to sort microglia (Control Sham and Control + WBRT, CD45^low/int^CD11b^+^) and repopulated cells (CSF-1R inhibitor Sham and CSF-1R +WBRT, CD45^low/int^CD11b^+^) for RNA extraction and RNA sequencing (Figure 1a).

**Figure 1:**
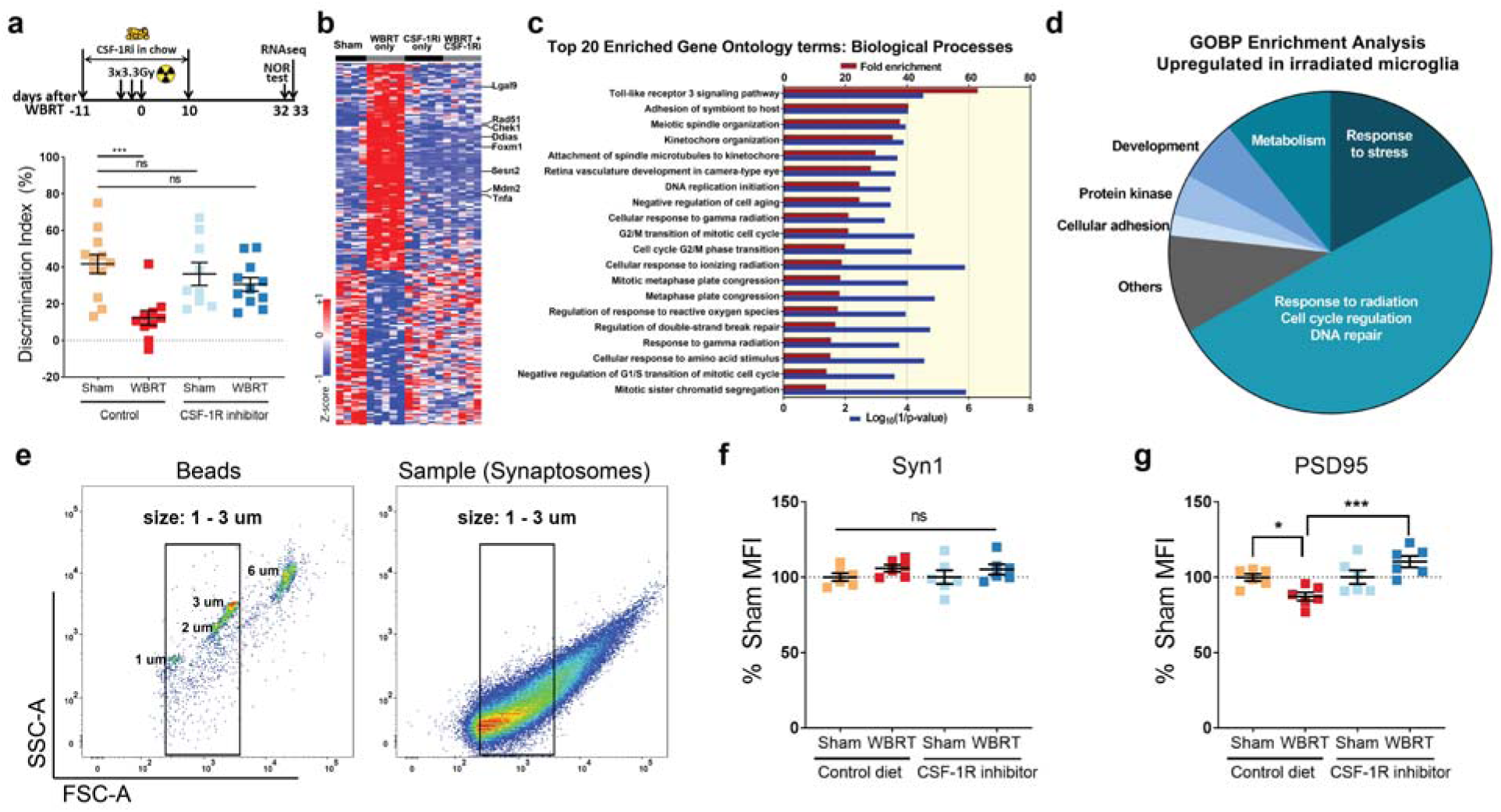
Microglia depletion and repopulation prevents long term radiation-induced memory deficits and loss of hippocampal PSD95. **a** experimental design and Novel Object Recognition (NOR) test result. CSF-1R inhibitor was used to deplete microglia during 3 doses of 3.3 Gy of whole-brain radiotherapy (WBRT). A 4-day NOR protocol was used to measure recognition memory, which ended on day 32 post WBRT. Microglia were isolated using fluorescent activated cell sorting (FACS) on day 33. and dot plots showing NOR results. Statistical analysis was performed using two-way ANOVA with Dunnett’s multiple comparisons test. There is no CSF-1Ri treatment effect (F(1,38)=1.787, p=0.1893), but significant WBRT effect (F(1, 38)=13.23, p=0.0008) and interaction between CSF-1Ri treatment and WBRT (F(1,38)=6.07, p=0.0184), N = 9-12, animals with insufficient exploration time on NOR test day were excluded. **b** hierarchically clustered heatmap showing significantly altered microglial genes by WBRT, but not changed with CSF-1Ri treatment. **c** bar graphs summarizing fold enrichment and p values of the top 20 enriched Biological Processes by Gene Ontology analysis from up-regulated microglial genes after WBRT (full list in Supplementary Table1). No significantly enriched terms were identified by GO analysis from down-regulated genes by WBRT. **d** a pie chart summarizing all enriched GOBP terms. ns= not significant, ***p<0.0001. **e** scatter plots showing gating strategy in flowsynaptometry analyses. Fluorescent beads at various sizes were used as standard to gate isolated hippocampal cell membrane fractions. Particles between 1 µm and 3 µm were considered synaptosomes and used to determine Synapsin1 and PSD95 protein levels by mean fluorescent intensities (MFIs). **f** dot plots to compare Synapsin1 and PSD95 MFI levels in hippocampal cell fractions. Statistical analyses were performed using two-way ANOVA with Tukey’s multiple comparisons test. ns = not significant, *p<0.05, ***p<0.001. N=6.

### Microglia depletion and repopulation eliminates radiation-induced transcriptome signatures

To identify biological pathways involved in radiation-induced memory deficits, we listed genes differentially expressed in microglia after WBRT with and without microglia depletion and repopulation for Gene Ontology Biological Process (GOBP) enrichment analysis. 204 genes were found to be differentially expressed (DE genes) only in microglia isolated from irradiated brains (Figure 1b **and Supplementary Table 1**). No enriched GOBP terms were found from the 87 WBRT down-regulated genes (Supplementary Table 1). There were 193 enriched GOBP terms from the 117 WBRT up-regulated genes, the top 20 enriched GOBP terms are listed in Figure 1c. Almost half (96) of these enriched GOBP terms were associated with increased response to cell cycle regulation, radiation, DNA repair and stress; the rest enriched GOBP terms were associated with increased metabolism (21), development (12), regulation of protein kinase activity (8), cellular adhesion (4) and other functions (Figure 1d, Supplementary Table 1). Notably, regardless of WBRT, the expression of these WBRT-induced DE genes did not change in cells isolated from brains treated with CSF-1Ri. These results demonstrate that the transcriptomic changes in microglia induced by WBRT can be completely eliminated after microglia depletion and repopulation.

To validate the RNAseq results, we next performed qPCR analyses using sorted microglia from animals in the same cohort (Figure 1b, and **Supplementary Table 1**). The expression of the toll-like receptor 3 (TLR3) family gene *Lgals9* was significantly increased by irradiation (WBRT + Control diet versus Sham + Control diet) and was at levels comparable to the shams (Sham + Control diet) when treated with CSF-1Ri despite of irradiation (Supplementary Figure S1a). *TNFα*, another TLR3 family member which also belongs to GOBP “regulation of response to reactive oxygen species (ROS)”, was significantly upregulated by irradiation (WBRT + Control diet versus Sham + Control diet); its expression levels are comparable between the Sham + Control diet and the WBRT + CSF1Ri treated groups. However, *TNFα* remained elevated in microglia from mice treated only by CSF-1Ri (Supplementary Figure S1b). Another gene from the GOBP “regulation of response to ROS”, *Sesn2*, was also significantly upregulated by WBRT (Supplementary Figure S1c). *Sesn2* remained at the control sham levels in CSF-1Ri only group and was significantly down-regulated in the WBRT + CSF-1Ri group. *Mdm2*, a gene that belongs to GOBP “cellular response to ionizing radiation”, was increased after WBRT, and significantly downregulated in in CSF-1Ri treated groups (Supplementary Figure S1d). Other WBRT-induced expression of radiation induced genes, *Ddias*, *Rad51, FoxM1* and *Check 1*, were all at the control sham levels in repopulated microglia regardless of the exposure to WBRT (Supplementary Figure S1 e – h). In conclusion, the qPCR validation confirmed that the transcriptomic changes seen in our RNAseq dataset were reliable. These results suggest that CSF-1Ri mediated microglia depletion during WBRT followed by repopulation eliminated radiation-induced signatures in the microglia transcriptome.

### Microglia depletion and repopulation prevents radiation-induced loss of hippocampal PSD-95

We previously demonstrated that brain irradiation resulted in reduced density of dendritic spines in hippocampal neurons (13). To accurately determine the effect of WBRT in the intrinsic synaptic protein levels we measured pre- (Syn1) and post-synaptic (PSD-95) markers in the hippocampus by flow-synaptometry (26, 27). Fractionated hippocampal cell membranes containing synaptosomes were enriched and particles between 1 – 3 µm were analyzed to measure synaptic protein levels using mean fluorescent intensities by FACS (Figure 1e). We observed no changes in pre-synaptic Synapsin-1 protein levels in the hippocampi across all groups (Figure 1f). However, we measured a significant reduction in post-synaptic protein PSD-95 after WBRT, which was completely prevented by CSF-1R inhibitor mediated microglia depletion (Figure 1g). These results cement the role of microglia in the radiation-induced loss of post-synaptic components after WBRT.

### Repopulated microglia after WBRT originate from peripheral monocytes

The fractalkine receptor CX3CR1 is expressed in both microglia and peripheral monocytes (28), while chemokine receptor CCR2 is mainly expressed in monocytes (29). In the Cx3cr1^GFP/+^Ccr2^RFP/+^ reporter mice, the different expression patterns of GFP and RFP can be used to distinguish microglia (GFP+RFP-) from monocytes (GFP+RFP+) (29). To investigate the cell-of-origin of repopulated microglia in our experimental paradigm we generated bone marrow chimeras with head-protected irradiation using fluorescent labeled bone marrow from Cx3cr1^GFP/+^Ccr2^RFP/+^ donor mice (Figure 2a). This allowed partial replacement of bone marrow cells without changing the permeability of the blood-brain-barrier (2, 30, 31). At 6 weeks after bone marrow transplantation about two thirds of peripheral monocytes were replaced by transplanted cells with fluorescent labels (Figure 2b). Bone marrow chimera animals were then treated with WBRT and CSF-1R inhibitor following the same experimental timeline used for RNA sequencing (Figure 2a). Next, we compared the compositions of myeloid cells in the brain after CSF-1R inhibitor-mediated depletion and repopulation. Flow cytometry analyses performed 33 days after WBRT revealed that microglia depletion and repopulation alone (Sham + CSF-1Ri) only resulted in limited accumulation of transplanted cells in the brain (Figure 2c**, BMT only**). However, in mice that received WBRT and CSF-1R inhibitor two thirds of the microglia were replaced by Cx3cr1-GFP labeled cells, close to the chimera efficiency (Figure 2 b **and** c**, BMT +fWBI**). These results suggest that microglia depletion during WBRT resulted in significant contribution of the CNS microglia pool by peripheral monocyte-derived BEMs.

**Figure 2:**
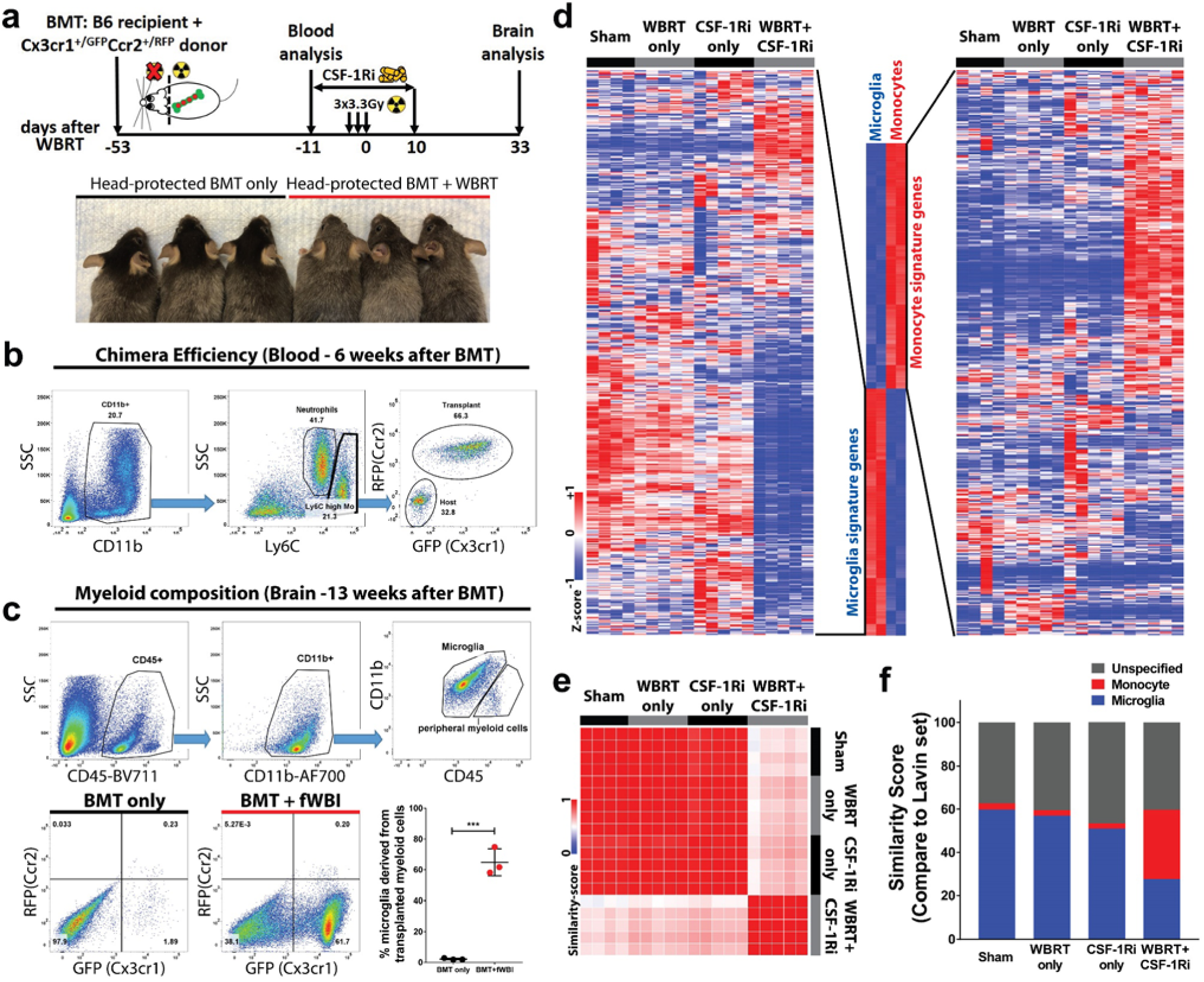
Repopulated microglia-like cells after depletion and WBRT originate from peripheral monocytes and retain monocytic signatures. **a** experimental design of head-protected bone marrow transplantation (BMT) followed by CSF-1Ri-mediated microglia depletion and WBRT. Lower panel shows fur colors before euthanasian for brain analysis. **b** representative FACS analysis gating strategy to analyze bone marrow chimera efficiency 6 weeks after BMT, about two thirds of the CD11b^+^Ly6C^high^ monocytes are replaced by GFP^+^RFP^+^ cells derived from donor bone marrow cells. **c** representative FACS analysis gating strategy and brain myeloid composition results. Upper panel shows FACS gating using CD45 and CD11b staining; microglia and microglia-like cells are defined by positive CD11b staining and low or intermediate CD45 levels. Lower panel shows scatter plots of GFP/RFP fluorescent levels of the CD11b^+^CD45^low/intermediate^ population in the brain, and a dot plot comparing percentages of peripheral myeloid cell derived microglia-like cells. Statistical analysis was performed using unpaired t-test, ***p<0.001. **d** hierarchically clustered heatmaps to compare microglia and monocyte signatures. A signature gene list was defined using a dataset published by Lavin and Winter et al, GSE63340. Defined list and expression details are in Supplementary Table 2). **e** Similarity matrix comparisons using defined monocyte and microglia signature genes. **f** bar graph showing similarity scores to compare relative numbers of genes (in percentage of the defined list) that express in the same trends as monocytes or microglia based on the Lavin and Winter *et al* dataset.

### Brain-engrafted macrophages retain monocyte signatures

We next assessed the transcriptomic profile of the BEMs after microglia depletion and WBRT by comparing our RNAseq dataset with a previous report by Lavin and Winter *et al* (32). To minimize false discovery and noise signals, we examined 1201 genes from this published dataset with a fold change greater than 1.50 or smaller than 0.667 for down-regulated genes (FDR <0.01, monocyte compared to naïve microglia), and found that 1066 genes were expressed in our samples (**Supplementary Table 2**). Strikingly, the hierarchical clustering of 525 monocyte- and 541 microglia-signature genes revealed that the expression profile of monocyte-derived BEMs (WBRT + CSF-1Ri) does not cluster with naïve (Sham + Control diet), irradiated (WBRT + Control diet) or repopulated (Sham + CSF-1Ri) microglia (Figure 2d). Similarity matrix analysis using this microglia/monocyte signature gene list revealed that the expression pattern in BEMs is different from naïve, irradiated and repopulated microglia (Figure 2e). Next, we counted genes in each group that expressed in the same trends as microglia or monocyte signature genes from the Lavin data set to determine the similarity scores to these two cell populations. We found that naïve, irradiated and repopulated microglia had 60%, 57% and 51% (718, 685 and 612) genes expressed in the same trends as microglia signature genes, respectively, with minimum similarity (2-3%) to monocyte signature genes; while BEMs expressed both microglia (28%, 331 genes) and monocyte signature genes (32%, 386 genes) (Figure 2f).

To validate microglia and monocyte signature genes that were differentially expressed in our RNAseq results we performed qPCR analyses (Supplementary Figure 2 **and Supplementary Table 2**). Microglia signature genes *Sall1*, *P2ry12*, *Tmem119* and *Trem2* were expressed at comparable levels in naïve, irradiated and repopulated microglia, while at significantly lower level in BEMs (Supplementary Figure 2 a – d). On the other hand, expression of monocyte signature gene *Runx3*, was significantly higher in BEMs than other groups (Supplementary Figure 2 e). Notably, previously reported brain-engrafted macrophage specific genes *Lpar6* and *Pmepa1* (*33*) have significantly higher expression levels in BEMs after CSF-1Ri and WBRT treatments compared to other groups (Supplementary Figure 2 f **and** g). In addition, the expression of *Ccr2*, a monocyte signature gene that was not differentially expressed in our RNAseq dataset, was also not differentially expressed among the four experimental groups by qPCR, suggesting that monocyte derived BEMs had lost their Ccr2 expression at this time point (Supplementary Figure 2 h). Taken together, these results confirm that after WBRT and CSF-1R inhibitor-mediated microglia depletion BEMs originate from peripheral monocytes.

### Monocyte-derived brain-engrafted macrophages resemble embryonic microglia signatures

Because monocyte-derived BEMs were exposed to the brain microenvironment for a short period of time, we hypothesized that they were functionally immature. To test this hypothesis, we first examined genes that were highly expressed at different developmental stages in microglia, and used yolk sac/embryonic and adult-specific genes as references (called embryonic and adult signature genes hereon) (34). Hierarchical clustering of 1617 embryonic and 785 adult microglia signature genes revealed that transcriptomes of BEMs were highly similar to embryonic microglia, while the transcriptomes of microglia from other groups were similar to adult microglia and did not resemble the embryonic microglia (Supplementary Figure S3a**, and** Supplementary table 3). In addition, a similarity matrix analysis using all 2402 overlapped genes between two datasets showed that BEMs had the lowest similarity with microglia from other groups (Supplementary Figure S3b). In addition, 54% of the listed genes (n=1306) in BEMs expressed in the same trends as *yolk sac*/embryonic microglia compared to adult microglia (Supplementary Figure S3c). In contrast, naïve (Sham), irradiated (WBRT only) and repopulated microglia (CSF-1Ri only) had much lower embryonic signature similarity scores (16%, 19% and 17%, n=381, 445 and 405, respectively, Supplementary Figure S3c). Notably, naïve, irradiated and repopulated microglia transcriptomes had high adult signature similarity scores (69%, 59% and 63%, n=1649, 1409 and 1507, respectively), while BEMs had the lowest adult similarity score (32%, n=759). These data suggest that the monocyte-derived BEMs start to resemble microglia by expressing more embryonic than adult microglia transcriptomic signature genes.

### Radiation-induced aberrant phagocytosis activity is abrogated in brain-engrafted macrophages

Aberrant loss of synapses during neuroinflammatory conditions has been linked with increased engulfment of synaptic compartments by microglia (35). To determine if WBRT affects the phagocytosis potency of microglia, we injected pre-labeled synaptosomes from a naïve donor mouse into the hippocampi of mice after WBRT and CSF-1R inhibitor treatment and measured engulfment by microglia using flow cytometry (Figure 3a). After WBRT there was a significant increase in the number of microglia engulfing synaptosomes in the hippocampus compared to naïve non-irradiated animals (Figure 3b). Strikingly, synapse engulfment activity was unchanged compared to naïve animals in animals treated with CSF-1R inhibitor during WBRT (Figure 3b). Immunofluorescent imaging at the injection sites confirmed that the injected synaptosomes were indeed engulfed by microglia, and the increased trend of engulfment by irradiated microglia remained unchanged (Figures 3 c **and** d, Supplementary Figure 4a). Notably, after hippocampal injection of fluorescent labeled latex beads into the hippocampus, we found that WBRT resulted in increased engulfment of latex beads was also inhibited by CSF-1R inhibitor treatment, suggesting that the WBRT-induced increase of engulfment was not specific to synaptosomes, but rather a general increase of phagocytosis potency (Supplementary Figure 4b). These data are the first to demonstrate that WBRT results in an increase in microglial phagocytosis activity in the hippocampus that can be prevented by transient microglia depletion and full repopulation.

**Figure 3:**
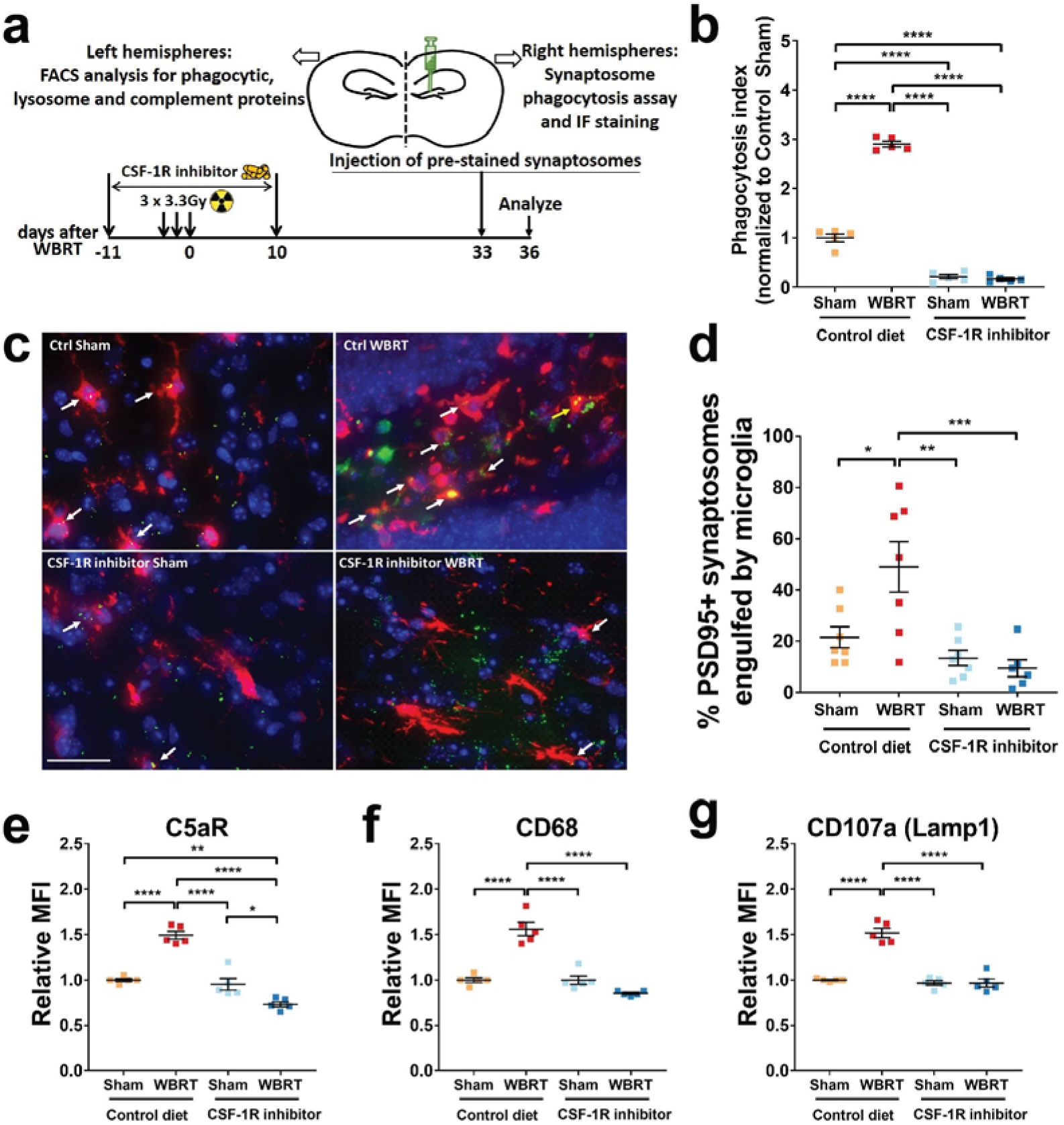
Repopulated microglia and brain-engrafted macrophages are not activated and phagocyte less synaptic compartments. **a** experimental design for in vivo synaptosome phagocytosis assays. Injection of pre-stained synaptosomes was timed to be the same as previous experiments. Three days later, on day 36 after WBRT, ipsilateral hemispheres were harvested and used for engulfment measurement using FACS or Immunofluorescent staining. **b** FACS analysis result showing levels of microglia that engulfed pre-stained PSD-95 signals. **c** representative images showing engulfment of pre-stained synaptosomes by microglia near injection site. White arrows point at microglia that have engulfed pre-stained synaptosomes. scale bar = 20 µm. **d** dot plot to show quantification result of synaptosome engulfment by immunofluorescent staining. **e – g** dot plots showing cell surface C5aR, and intracellular CD68 and CD107a protein levels in microglia and BEMs. Statistical analyses were performed using two-way ANOVA with Tukey’s multiple comparisons test. *p<0.05, **p<0.01, ***p<0.001, ****p<0.0001. N = 5 – 6.

### Irradiation-induced complement and phagocytic receptors expression in microglia are absent in BEM after WBRT

Microglial complement receptors play essential roles in physiologic synaptic elimination during development and aberrant elimination during neuroinflammatory conditions (35, 36). To understand the mechanisms of increased microglia phagocytic activity after WBRT, we measured expression levels of a list of complement receptors, phagocytic markers and lysosome proteins in microglia by flow cytometry. The expression of complement receptor C5aR was significantly elevated in microglia at one month after WBRT. However, in animals treated with CSF-1Ri C5aR expression was unchanged from naïve animals (Figure 3e). The same trend was observed in the expression levels of CD68 and lysosomal-associated membrane protein 1 (LAMP-1) (Figure 3 f **and** g). These results were consistent with our data demonstrating decreased PSD95 levels (Figure 3b) and increased microglial phagocytosis activity in the hippocampus after WBRT (Figure 3 b **and** d). In addition, complement receptor CR3 (CD11b) was significantly elevated in microglia after WBRT or CSF-1Ri treatments alone, and remained unchanged in BEMs with combined WBRT and CSF-1Ri treatments (Supplementary Figure S5a). No changes in the complement receptor C3ar1 were measured after WBRT or CSF-1R inhibitor treatment (Supplementary Figure S5b). These results demonstrate that the increased microglia phagocytosis of synaptosomes after WBRT was associated with increased phagocytic and lysosome proteins, and was likely through the complement pathways.

### Brain-engrafted macrophages after microglia depletion persist in the brain

To determine whether BEMs are present long-term in the brain, we monitored this cellular population for 6 months after WBRT. To eliminate the limitation of using bone marrow obtained from the Cx3cr1^+/GFP^Ccr2^+/RFP^ knock-in reporter mouse strain, we used an actin-GFP transgenic line as bone marrow donors and generated chimeras using the same body-only irradiation protocol (Figure 4 a). Six weeks later, mice were treated with CSF-1R inhibitor and WBRT and then used to trace BEMs at varies time points (Figure 4 a). Whole coronal sections at the level of the dorsal hippocampus were stained with Iba1 and imaged to quantify total Iba1+ and GFP+ cells (Supplementary Figure 6 a). We found that all GFP+ cells in the brain were also Iba1+, suggesting that BEMs were indeed derived from the periphery. In addition, the morphology of Iba1+GFP+ BEMs were analyzed and compared to Iba1+GFP-microglia (Figure 4 b **and** d). We found that round-shaped Iba1+GFP+ BEM cells entered the brain starting from 7 days after WBRT, and started to obtain more processes that resembled microglia morphology (Figure 4 b). However, Sholl analysis demonstrated that the morphology of BEMs remained stable from 33 days after WBRT and never reached the structural complexity of microglia (Figure 4c, Supplementary Figure S7). We found that 40 – 90% of Iba1+ cells are also GFP+ at 14 days after WBRT. This ratio remained at high levels at 1, 3 and 6 months after WBRT (Figure 4 e). Interestingly, although the Iba1+ and Iba1+GFP+ cell numbers are not fully recovered at 14 days after WBRT, the microglia replacement ratio was similar to the level of later time points (Figure 5 e and Supplementary Figure S6 b and c), suggesting a non-competitive manner of brain parenchyma occupancy by microglia and BEMs. These data demonstrate that BEMs enter the brain shortly after WBRT, adapt to a microglia-like morphology and maintain a stable population.

**Figure 4:**
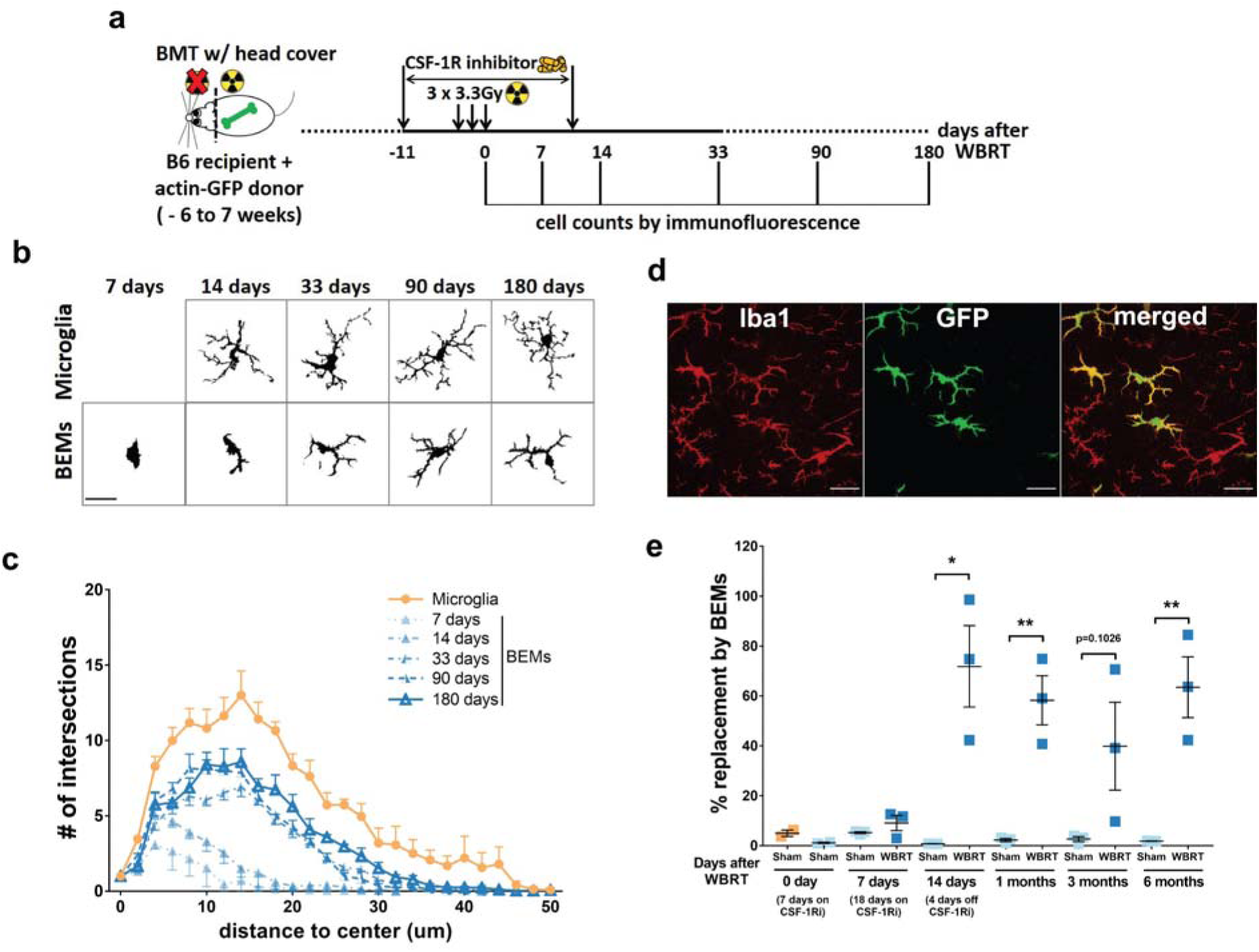
BEMs gradually adapt to microglia-like morphology and persist in the brain. **a,** schematic of experimental design for long-term assessment of BEMs. **b,** representative images of microglia/BEMs counting, scale bar = 20 μm. **c**, Sholl analysis results showing numbers of intersections at different distances to cell center, BEMs at 7, 14, 33, 90 and 180 days after WBRT were compared to naïve microglia age-matched to 90 days after WBRT., representative images showing differential Iba1 and GFP expressing bprofiles of microglia (Iba1+ GFP-) and BEMs (Iba1+ GFP+) in a BEM bearing brain at 33 days after WBRT. **e**, dot plot to show percentage of replacement of microglia by BEMs, each dot represent an individual mouse. n = 2 -3. Statistical analyses were performed using unpaired t-test at each distance point (c) or time point (e). See Supplementary Figure 7 for detailed comparisons between microglia and BEMs at each time point.

**Figure 5:**
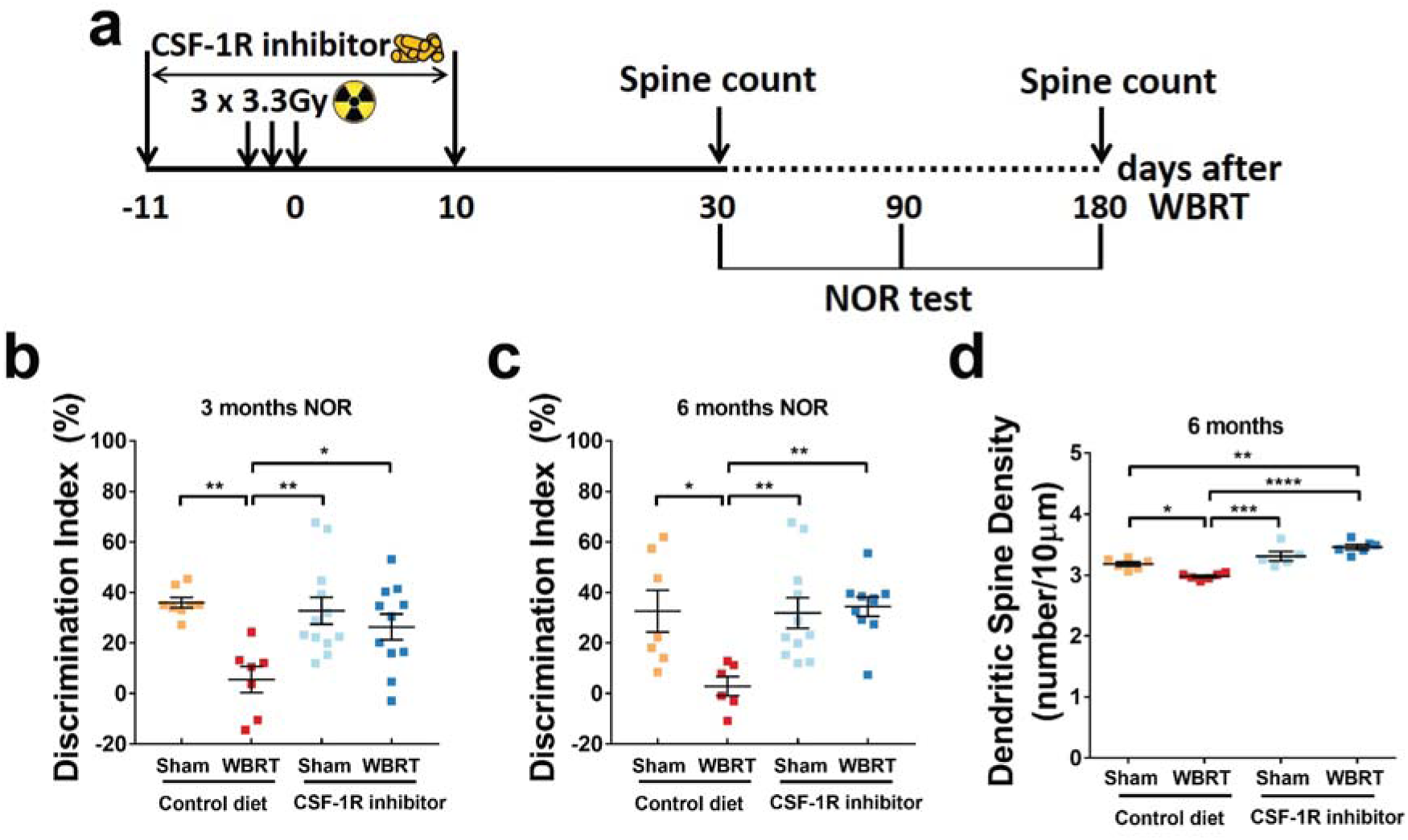
BEMs provide long-term protection against WBRT-induced dendritic spine and memory loss. **a** schematic of experimental design for long-term memory and dendritic spine density analyses. **b** and **c** dot plots to show NOR test results at 3 and 6 months after WBRT, respectively. N = 6–12. **d** dendritic spine counts of hippocampal granule neurons at 6 months after WBRT, N = 5 – 6. Statistical analyses were performed using two-way ANOVA with Tukey’s *post hoc* multiple comparisons test (**b** - **d**). *p<0.05, **p<0.01, ***p<0.001, ****p<0.0001.

### BEMs provide long-term protection against WBRT-induced memory deficits and hippocampal dendritic spine loss

To measure the long-term cognitive outcomes, we treated a batch of wildtype animals, and tested their recognition memory at 3 and 6 months after WBRT (Figure 5 a). We found that WBRT resulted in persistent loss of recognition memory also at 3 and 6 months, while CSF-1Ri treatment alone did not alter recognition memory performance (Figure 5 b **and** c). Strikingly, mice that received WBRT along with temporary microglia depletion did not show any memory deficits and performed undistinguishable from control animals at 3 and 6 months (Figure 5 b **and** c). Our previous report demonstrated that WBRT-induced dendritic spine loss in hippocampal neurons was fully prevented by temporary microglia depletion during irradiation (13). In this study, we sought to understand if the protective effects persisted up to 6 months after WBRT. Our results clearly show that radiation-induced loss of dendritic spines in hippocampal neurons persists to this time point, and that the protective effect of microglia depletion and subsequent replacement by BEMs is long lasting (Figure 5 d). Taken together, brief depletion of microglia during WBRT induces sustainable BEMs in the brain and provides long-term protection against irradiation-induced deficits in recognition memory.

### Replacement of microglia by BEMs protects against concussive injury-induced memory loss

To investigate the function of BEMs after they replaced microglia, a single mild concussive <u>C</u>losed <u>H</u>ead <u>I</u>njury (CHI) was given to mice 30 days after CSF-1R inhibitor treatment and WBRT; microglia/BEMs morphology and phagocytic activities were measured following recognition memory test by NOR (Fig 6 a). By FACS analysis at 24 days post injury, we found that phagocytosis activity increased (p=0.0419) after CHI in microglia but not in BEMs (Figure 6 b). Quantification of immunofluorescent staining of Iba-1/PSD-95 co-localization also revealed a trend of increased engulfment towards pre-stained synaptosomes by microglia but not by BEMs (Fig 6 c **and** d). In addition, the structural complexity of microglia decreased in Sholl analysis, while the morphology of BEMs remained unchanged after CHI (Figure 6 e). Furthermore, at 20 days after injury, CHI-induced recognition memory deficits were spared in mice whose microglia were replaced by BEMs (Figure 6 f). These results demonstrate that unlike resident microglia which transition to a less ramified morphology and exhibit increased phagocytic activity towards injected synaptosomes, BEMs remain unchanged in both morphology and phagocytic activity in response to CHI. More importantly, our data suggest that replacement of microglia by BEMs can protect against CHI-induced memory loss.

**Figure 6:**
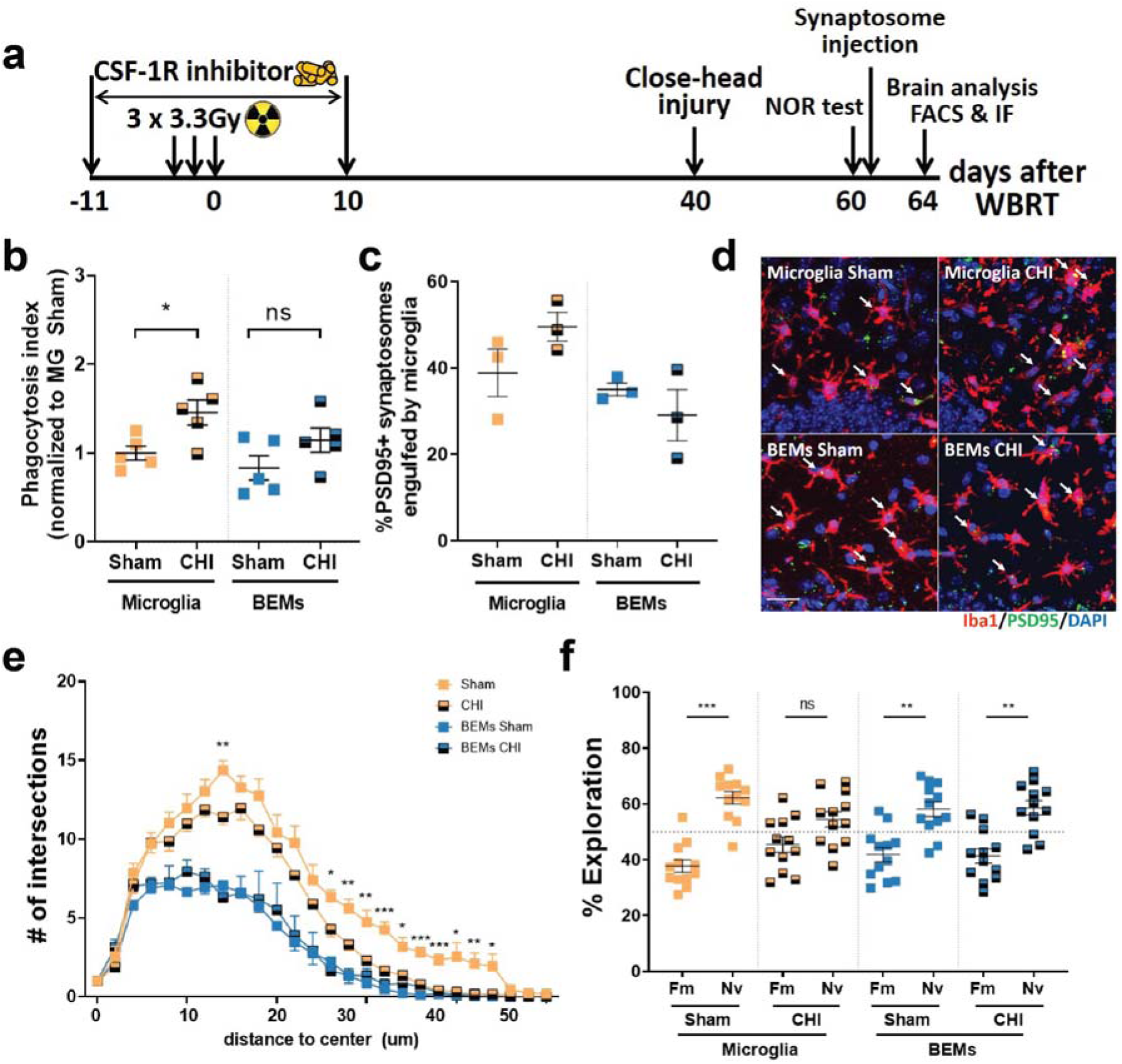
BEMs protects against concussive injury-induced memory deficits. a, schematic of experimental design for concussive injury, cognitive test and following analyses. b, dot plot to show the result of in vivo phagocytosis assay by FACS after injection of pre-stained synaptosomes, each dot represents value from an individual mouse, n = 4 -5. c, dot plot showing result of in vivo phagocytosis assay by IF imaging and quantification, each dot represents mean counts from an individual mouse, n = 3. d, representative images showing microglia and BEMs (arrows) phagocyting injected synaptosomes (green dots). e, Sholl analysis result showing numbers of intersections at different distances to the cell center, n = 5 - 6. f, dot plot showing NOR test result, each dot represent the performance of an individual mouse, n = 12. Statistical analyses were performed using two-way ANOVA with Tukey’s multiple comparisons test (b and c) for each distance point (e) or unpaired t-test (f). *p<0.05, **p<0.01, ***p<0.001.

## Discussion

Here we provide evidence for the direct involvement of microglia phagocytic activity towards synaptic compartments as a mechanistic cause for loss of dendritic spines with consequent impairments in memory functions after WBRT. Replacement of microglia with monocyte derived BEMs prevents loss of synapses and consequent memory deficits. Importantly, BEMs replacing microglia are also protective against a second injury to the brain. Together our results unravel novel immediate and long lasting therapeutic benefits of microglia depletion and repopulation during WBRT.

Microglia play pivotal roles in reshaping synaptic networks during neonatal brain development (37, 38). They engulf synaptic elements by active synaptic pruning in an activity- and complement-dependent manner (38). Microglia-driven aberrant loss of synapses and consequent impairment of cognitive functions have also been reported in animal models of AD (35), infection (39), injury (40, 41), and aging (42). Using RNA sequencing, we compared the transcriptomes of microglia from irradiated and non-irradiated brains after CSF-1Ri-mediated microglia depletion and repopulation. WBRT induces increased expression of genes that mainly belong to cell cycle regulation, DNA damage repair and stress-induced biological processes (Figure 1d). As a result, activated microglia have higher engulfing potential towards both intrinsic and extrinsic synaptic compartments (Figures 1 g, Figure 3 b - d). This view is further supported by the increased expression of endosome/lysosome proteins CD68 and CD107a with the complement receptors CR3 and C5ar1 measured in microglia chronically after WBRT (Figure 3 e **–** g**, and** Supplementary Figure 5). Notably, both endosome/lysosome proteins and complement receptor expressions were comparable to naïve microglia (sham + control diet) in BEMs (WBRT + CSF-1Ri) and repopulated microglia (CSF-1Ri only). These results suggest that the loss of hippocampal synapses after WBRT may be dependent on the activation of the alternative complement pathway. Interestingly, although BEMs are morphologically similar to adult microglia, they retain a transcriptomic signature similar to both circulating monocytes and embryonic microglia (Figure 2 **and** Supplementary Figure 3).

The decrease in post-synaptic protein PSD95 level in hippocampal synaptosomes is also paralleled with reductions in hippocampal dendritic spines (Figures 5 d **and Feng et al**(13)). However, pre-synaptic Synapsin 1 protein levels are not affected by WBRT or microglia depletion, suggesting that WBRT mainly induces loss of post-synaptic compartments. Interestingly, although the phagocytosis potency of repopulated microglia and BEMs are both low (Figures 3 b **and** d), microglia depletion and repopulation alone does not affect dendritic spine density (Figures 5 d **and** e). On the other hand, microglia replacement by BEMs results in increased dendritic spine density compared to those with radiation alone, and microglia depletion alone (13). Strikingly, the protective effect of microglia depletion during WBRT results in protected memory functions and extends to 3- and 6-months following irradiation (Figure 5 b **and** c). The dendritic spine density in mice that received WBRT and CSF-1Ri remained higher than those who only received CSF-1Ri (Figure 5 e) suggesting that in an non-reactivate state (evidenced by no changes in genes involved in cell cycle and radiation response, in microglial phagocytosis and lysosome proteins, and in phagocytosis activity towards injected synaptosomes and latex beads) of repopulated microglia and BEMs may have intrinsic differences in maintaining the homeostasis of dendritic spines, which appears to diminish over time.

In the CNS, microglia maintain a stable population by self-renewal in either a random manner or through clonal expansion (3, 43). CSF-1R inhibitor treatment alone results in acute depletion of up to 99% of CNS resident microglia, with repopulated microglia arising solely from the residual microglia and their progenitor cells that remain after treatment (8), (44). The repopulated microglia have transcriptional and functional profiles similar to naïve microglia (9). Peripheral macrophages can engraft into the brain but remain morphologically, transcriptionally and functionally different from CNS resident microglia (45, 46),. Under specific circumstances, monocytes entering the CNS can become microglia-like cells. This is most clearly demonstrated in experiments where lethal whole-body irradiation was followed by bone marrow transplantation with labeled monocytes (Ccr2^+^Ly6C^high^), resulting in accumulation of these cells in the brain (30). In neonatal mouse brains monocytes can enter the brain parenchyma without head irradiation and become microglia-like cells at a low frequency (47). In addition, chronic depletion of microglia without irradiation also results in myeloid cells entering the CNS and becoming BEMs (33). However, the roles of BEMs in cognitive functions are largely unknown. Here we report that concurrent microglia depletion and therapeutic brain irradiation causes peripheral monocytes to enter the brain parenchyma and become microglia-like BEMs. BEMs enter the brain at 14 days after the completion of brain irradiation, or 4 days after the CSF-1Ri withdrawal (Figure 4 e**, and** Supplementary Figure 6). Notably, although the ratio of BEMs was high at this time point the total number of Iba1 positive cells is not fully recovered (Supplementary Figure 6 b). This ratio remains at high levels in head-irradiated mice throughout the current study (Figure 5 e), suggesting that microglia depletion during WBRT results in sustainable replacement of microglia by BEMs. Importantly, this observation correlates with long-term protection against WBRT-induced loss of recognition memory and dendritic spines in hippocampal granule neurons (Figure 6 b **–** e).

In the clinic, cancer patients are unlikely to receive a second round of radiotherapy to the brain. Therefore, instead of introducing a second round of WBRT, after they occupied the brain we gave BEM-bearing mice CHI that causes memory deficits (48, 49), and further examined BEMs’ response to a single head trauma. Our data show that microglia had increased phagocytic potential to exogenous synaptosomes after CHI, while phagocytic activity of BEMs did not change and remained at a similar level as naïve microglia (Figure 6 b **and** c). This is further demonstrated by Sholl analysis of BEMs showing no change in morphology after CHI (Figure 6 e). Most importantly, CHI-induced memory deficit was prevented in BEM-bearing mice (**Figure 7 f**). These data are the first to demonstrate that BEMs can prevent brain injury-induced cognitive dysfunction.

A limitation of the current study is that we didn’t investigate transcriptomic profiles and phagocytic functions of BEMs at chronic time points after they replaced microglia. Therefore, it is unknown whether BEMs can obtain transcriptomic profiles and functions closer to adult microglia at later time points. In addition, microglia have been shown to mediate forgetting through complement-dependent synaptic elimination (50). We did observe increased dendritic spines in hippocampal granule neurons at both 1 and 6 months after WBRT compare to sham animals, suggesting that BEMs may have lower activity in eliminating synapses than naïve microglia. Therefore, the consequences of microglia replacement by BEMs in normal forgetting need further investigation. In addition, transcriptomic and functional studies at chronic time points with other microglia depletion models will provide insights into the transcriptomic and functional dynamics of BEMs in the brain.

In conclusion we report evidence for the mechanism by which microglia depletion and repopulation after WRBT prevents memory loss. Our data demonstrate that replacement of CNS resident microglia by peripheral monocyte-derived BEMs results in a transcriptional and functional reset of immune cells in the brain to an inactive state, which spares the brain from WBRT-induced dendritic spine loss in hippocampal neurons and recognition memory deficits. Most importantly, replacement of microglia by BEMs protects against concussive brain injury-induced cognitive deficits. These results suggest that replacement of depleted microglia pool by peripheral monocyte-derived BEMs represents a potent treatment for irradiation-induced memory deficits.

## Materials and Methods

### Animals

All experiments were conducted in compliance with protocols approved by the Institutional Animal Care and Use Committee at the University of California, San Francisco (UCSF), following the National Institutes of Health Guidelines for Animal Care. 7 weeks old C57BL/6J male mice were purchased from the Jackson Laboratory and housed at UCSF animal facilities and were provided with food and water ad libitum. All mice were habituated for one week before any treatments or procedures. 8–10 weeks old Cx3cr1^GFP/+^Ccr2^RFP/+^ mice were breed by crossing the Cx3cr1^GFP/GFP^Ccr2^RFP/RFP^ line with wildtype C57BL/6J mice, and used as donors for the bone marrow chimeras.

### CSF-1Ri treatment

CSF-1Ri (PLX5622 formulated in AIN-76A standard chow at 1200 ppm, Research Diets, Inc) were provided by Plexxikon, Inc (Berkeley, CA). Mice were given free access to either CSF-1Ri chow or control diet (AIN-76A without PLX5622) for 21 days. Approximately 4.8 mg of PLX5622 was ingested by each mouse per day in the treated group (calculation based on 4 g/mouse daily consumption).

### Fractionated whole-brain radiotherapy (WBRT)

8 weeks old mice were injected with ketamine (90mg/kg) /Xylazine (10 mg/kg) mix. When fully immobilized mice were placed in irradiator with cesium-137 source at the dose rate of 2.58 Gy/min. The body was shielded with a lead collimator that limited the radiation beam to a width of 1 cm to cover the brain. Three radiation fractions (3.3 Gy) were delivered every other day over 5 days. Sham animals received ketamine/xylazine without irradiation.

### Bone marrow chimeras

8 weeks old C57BL/6J mice were used as bone marrow recipients. 8 weeks old males received two doses of 6 Gy cersium-137 irradiation at the dose rate of 2.58 Gy/min with head protected by lead plates 6 hours apart. Bone marrow cells from 6–10 weeks old donors Cx3cr1^+/GFP^Ccr2^+/RFP^ or B6-EGFP (The Jackson Laboratory, stock No 003291) were isolated and resuspended in sterile saline at a concentration of 100 million cells/ml. 0.1 ml of bone marrow cells were injected into recipients via retro-orbital injection immediately after the second head protected irradiation. Bone marrow chimeras were housed with 1.1 mg/ml neomycin as drinking water for 4 weeks and allowed an additional 2 weeks to recover before any treatments.

### Concussive TBI – Closed head injury

12 weeks old C57BL/6J mice were randomly assigned to each TBI or sham surgery group. Animals were anesthetized and maintained at 2-2.5% isoflurane during CHI or sham surgery. Animals were secured to a stereotaxic frame with nontraumatic ear bars and the head of the animal was supported with foam. Contusion was induced using a 5-mm tip attached to an electromagnetic impactor (Leica) at the following coordinates: anteroposterior, −1.50 mm and mediolateral, 0 mm with respect to bregma. The contusion was produced with an impact depth of 1 mm from the surface of the skull with a velocity of 5.0 m/s sustained for 300 ms. Animals that had a fractured skull after injury were excluded from the study. Sham animals were secured to a stereotaxic frame with nontraumatic ear bars and received the midline skin incision but no impact. After CHI or sham surgery, the scalp was sutured and the animal was allowed to recover in an incubation chamber set to 37 °C. All animals recovered from the surgical procedures as exhibited by normal behavior and weight maintenance monitored throughout the duration of the experiments.

### Synaptosome isolation staining and injection

Fresh hippocampi from a naïve mouse was homogenized and spun down in 0.32M sucrose solution (dissolved in 50 mM HEPES buffer). Supernatant was centrifuged in 0.65M sucrose solution at 12,000 rpm for 30 minutes at 4°C. The synaptosome containing pellet was resuspended in 1 x ice-cold PBS, diluted to 100 µg/ml, and stained with PSD-95 antibody (Millipore) on ice for 30 minutes followed by a secondary antibody staining (Invitrogen, goat anti-mouse AF488). Stained synaptosomes were washed and diluted 20 times in PBS and stored at -80°C. 2 µl of pre-stained synaptosomes were injected into the right hippocampus of each recipient mouse at the coordinate relative to the bregma: AP + 1.6 mm, ML + 1.6 mm and DV -2.0 mm. Mice were euthanized 3 days later. The left hemispheres (uninjected) were used for phagocytosis markers staining and the right hemispheres (injected) were used to assess synaptosome phagocytosis levels by flow cytometry or immunofluorescent staining.

### Immunofluorescent Staining

hemi-brains with synaptosome injection were fixed in 4% PFA overnight, cryo-protected in 30% sucrose solution in 1 x PBS and sliced in 20 µm sections. Sliced tissues were stained with Iba1 (Fujifilm Wako Pure Chemical Corporation, 019-19741) followed by a secondary antibody staining (goat anti-rabbit AF568, Invitrogen, A-11011). DAPI was used for nuclear staining. Images close to the injection site (Supplementary Figure 1a) were taken using a Zeiss Imager Z1 microscope under a 20x objective lens. Tissues from bone marrow chimeras were processed and stained as described above. Images were taken using a CSU-W1 Nikon Spinning Disk Confocal microscope under 10x air, 20x air or 100x immerse oil lenses. All images were analyzed using the Fiji/ImageJ software by experimenters blinded to sample information.

### Behavior test

Novel Object Recognition (NOR) task was used to test hippocampal dependent recognition memory at one, three and six months after the last dose of irradiation. All tests took place during the dark cycle in a room with dim red light as previously described (13, 14). Briefly, mice were habituated in an open arena (30 cm x 30 cm x 30 cm, L x W x H) for 10 minutes on day one and day two. On day three, two identical objects were put into the arena at a distance of 21 cm and mice were allowed to explore for 5 minutes. On day four, one object was replaced by a novel object and mice were allowed to explore for 5 minutes. All trials were recorded by an overhead camera and analyzed using Ethovision software. Data are presented as discrimination Index, calculated using fomular DI = (T_Novel_ - T_Familiar_)/(T_Novel_ + T_Familiar_).

### Flow cytometry

mice were perfused with cold PBS after euthanasia. Brains were immediately removed and dissociated using a Neural Tissue Dissociation kit (P) (Miltenyi Biotec). Brain cells were resuspended in 30% Percoll solution diluted in RPMI medium, and centrifuged at 800 g for 30 minutes at 4°C. Cell pellets were washed with FACS buffer (1 x DPBS with 0.5% BSA fraction V and 2% FBS), blocked with mouse CD16/32 Fc block (BD Biosciences #553141) and stained with fluorophore conjugated antibodies (CD11b-AF700, CD45-FITC, BD Pharmingen 557690 and 553080, C5aR-PE, CD68-PE and CD107a-PE, Miltenyi Biotec 130-106-174, 130-102-923 and 130-102-219), washed with FACS buffer and used for sort or analyses of bone marrow chimera efficiency. Data were collected on an Aria III sorter using the FACSDIVA software (BD Biosciences, V8.0.1), and analyzed with Flowjo software (FlowJo, LLC, V10.4.2).

### Flow synaptometry

after isolation (described above) synaptosomes were stained with PSD-95 (Abcam ab13552) or Synapsin-1 (Millipore #1543) antibodies on ice for 30 minutes, washed and followed by a secondary antibody staining (Invitrogen, goat anti-mouse AF488, A-11001). Stained synaptosomes were used immediately for analysis of mean fluorescent intensity measurement. Fluorescent latex beads of 1 µm, 2 µm, 3 µm and 6 µm were used as references of particle sizes in the FSC-A vs SSC-A dot plot. Events between 1 µm and 3 µm were used to measure mean fluorescent intensities of isolated synaptosomes under the FITC channel. Data were collected on an Aria III sorter using the FACSDIVA software, and analyzed with Flowjo software. At least 100,000 events were collected from each sample for the analyses.

### RNA sequencing

mRNA was isolated from 100,000 to 400,000 sorted microglia or BEMs (CD45^low/int^/CD11b^+^ cells) using the Dynabeads mRNA DIRECT Purification Kit (Invitrogen #61011) following the manufacturer’s instructions. RNA sequencing libraries were generated using the Ovation RNA-seq system V2 and Ultralow Library Construction System sample prep kits (NuGEN). Libraries were sequenced on the HiSeq 2500 to generate single end 50bp reads according to the manufacturer’s instructions. Normalized per-gene read counts were used to compare relative gene expression levels across samples. Only genes with average read counts greater than 10 were included for analyses. Heatmaps were drawn using the online analysis software Morpheus (Broad Institute, https://software.broadinstitute.org/morpheus), followed by hierarchical clustering using the One minus pearson correlation method. Gene Ontology analysis was performed using the Statistical overrepresentation test (GO biological process complete, PANTHER version 14) ^(51)^. Bar graphs to visualize fold enrichment and p values of enriched GO biological pathways were drawn using the GraphPad Prism software (V 7.01, GraphPad Software, Inc). For analysis of monocyte/microglia signature genes, dataset from Lavin and Winter et al was used as reference (GSE63340) ^(32)^. Genes significantly up or down regulated (p<0.05, fold-change > 1.5 or <0.667) in monocytes vs microglia comparisons are defined as monocyte or microglia signature genes, respectively. Heatmaps were drawn as described above, and similarity matrix were drawn using the Morpheus online tool with Pearson correlation. Monocyte/microglia similarity scores were calculated based on the numbers of genes in each treatment group from this study that expressed in the same trend as monocyte/microglia signature genes (genes with fold-change between 0.6667 and 1.500 or with FDR>0.01 were defined as unspecified). For juvenile/embryonic signature analysis, dataset from Matcovitch-Natan and Winter et al was used as reference (GSE79819) ^(34)^. Gene listed to be highly expressed in Yolk Sac and embryonic day 10.5–12.5 were defined as embryonic/juvenile microglia signatures, genes highly expressed in adult cortex/hippocampus/spinal cord were defined as adult microglia signatures. Heatmaps, similarity matrix and similarity scores were drawn or calculated as described above.

### qPCR

mRNAs were extracted from sorted microglia using the Dynabeads mRNA DIRECT Purification Kit (Invitrogen #61011), and reverse transcribed into cDNAs using reverse transcription kit (info) . qPCR reactions were set up in duplicate reactions using the PowerUp SYBR Green Master Mix kit (Applied Biosystems #A25777) using an Mx3000P qPCR System (Agilent, Santa Clara, CA) following the manufacturer’s instructions. Data were analyzed using the standard curve method. Standard cDNAs were generated with total RNAs from mixed naïve and irradiated mouse brains. qPCR primers sequences are listed in Supplementary Table 4.

### Sholl analysis

Images of GFP+ (BEMs from bone marrow chimeras) or Iba1+ (AF555, all microglia cells, and BEMs from non-bone marrow chimeras) cells were acquired from stained frozen sections (20um) using a confocal microscope under 100x objectives (CSU-W1 Spinning Disk/High Speed Widefield, Nikon). Max Z-projections from Z-stack images (0.26um step size) were used for Sholl analysis(52) in Fiji(53) software using the following settings: manually defined cell center at the cell body, the numbers of intersections between cellular processes and circles with incremental radius (2um step size, up to 60um) were recorded, plotted and compared across samples.

### Statistical analyses

Two-way ANOVA was used to determine radiation and CSF-1Ri treatment effects for NOR, qPCR, flowsynaptometry, flow cytometry, immunofluorescent staining counts and dendritic spine count results, with Tukey’s post hoc multiple comparisons. One-way ANOVA with Sidak’s post hoc multiple comparisons was used to determine effect of developmental stages for dataset published by Matcovitch-Natan and Winter et al. Unpaired t-test was used to determine differentially expressed microglia/monocyte signature genes from dataset published by Lavin and Winter et al. Unpaired t-test was used calculate the p value of the comparison of BEMs contributions between the BMT and BMT + WBRT groups. Exact p values and numbers of animals used in each experiment were listed in each related figure legend. All error bars represent mean ± SEM.

## Acknowledgements

We thank the Nikon Imaging Center at UCSF for assistance with confocal microscope. We thank the Laboratory for Cell Analysis at UCSF for assistance with FACS sort and analysis.

## Author contributions

X.F. conceptualized the study, designed and performed the experiments, analyzed the data and wrote the manuscript. E.F. performed experiments, analyzed the data and wrote the manuscript. M.P performed experiments, analyzed the data and revised the manuscript. D.C. assisted in the in vivo phagocytosis assay and data analysis. Z.B. helped with the in vivo phagocytosis assay imaging and data analysis. M.B. assisted in the Sholl analysis of microglia and BEMs. S.G analyzed the long-term dendritic spine counts data. S.L. provided assistance in experiments related to WBRT and BM chimeras. N.G. provided critical inputs to the study and revised the manuscript. S.R. conceptualized and supervised the study and revised the manuscript. All authors approved the final version of the manuscript.

## Supplementary Figures

**Supplementary Figure 1:**
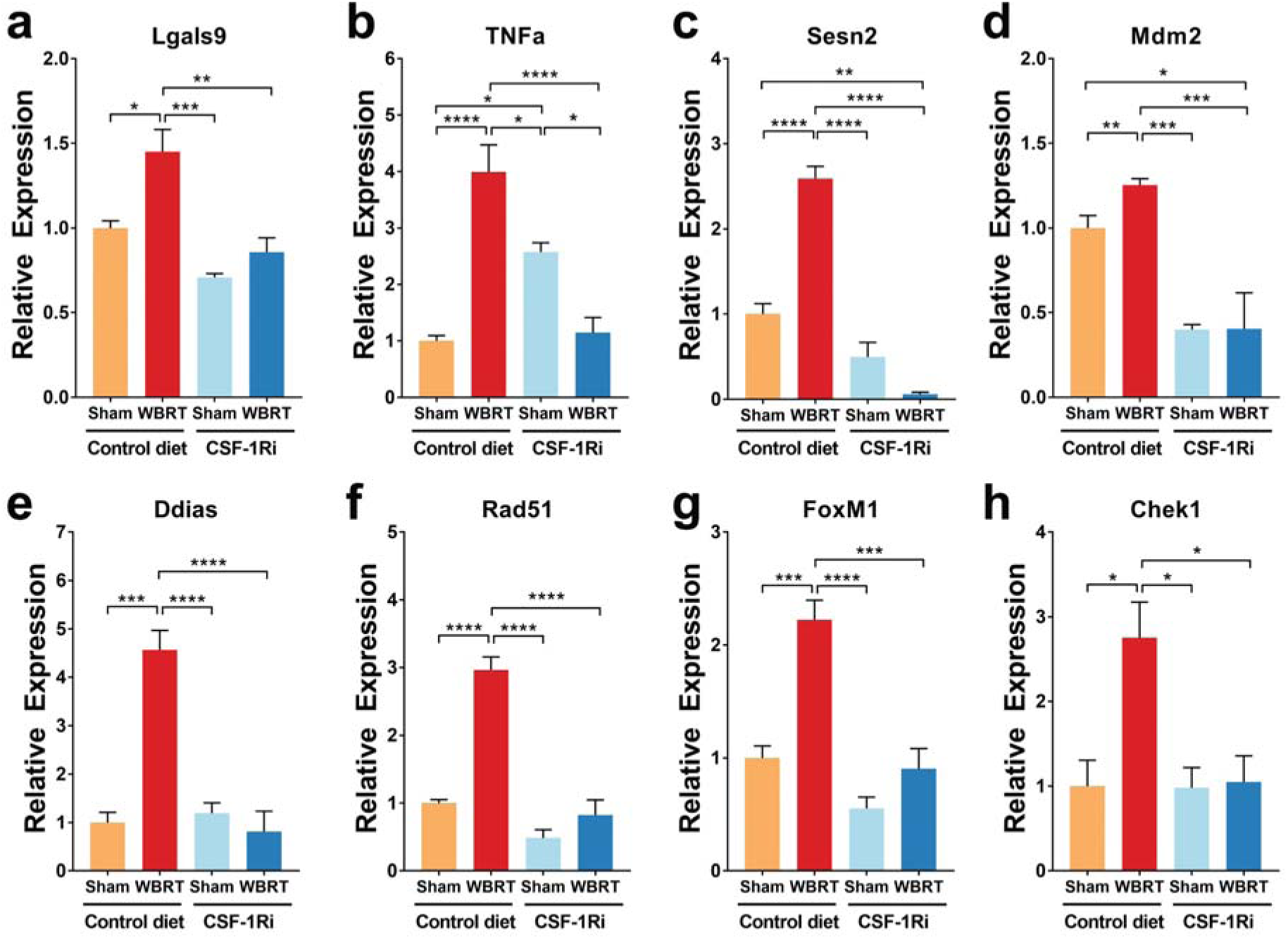
qPCR validation of radiation-induced genes. Genes from highly enriched GOBP terms were selected to validate RNAseq results. **a** and **b** Toll-like receptor 3 signaling pathway: *Lgals9* and *TNFα*. **c-e** Cellular response to ionizing radiation: *Rad51*, *Mdm2* and *Ddias*. **d** and **f** Regulation of response to reactive oxygen species: *TNFα* and *Sesn2*. **c, g and h** Regulation of double-strand break repair: *Rad51*, *Foxm1* and *Chek1*. Statistical analyses were performed using two-way ANOVA with Tukey’s multiple comparisons test. *p<0.05, **p<0.01, ***p<0.001, ****p<0.0001. N = 4–6. The qPCR experiments were performed in duplicates with similar results. Figures shown here are representative results from one experiment.

**Supplementary Figure 2:**
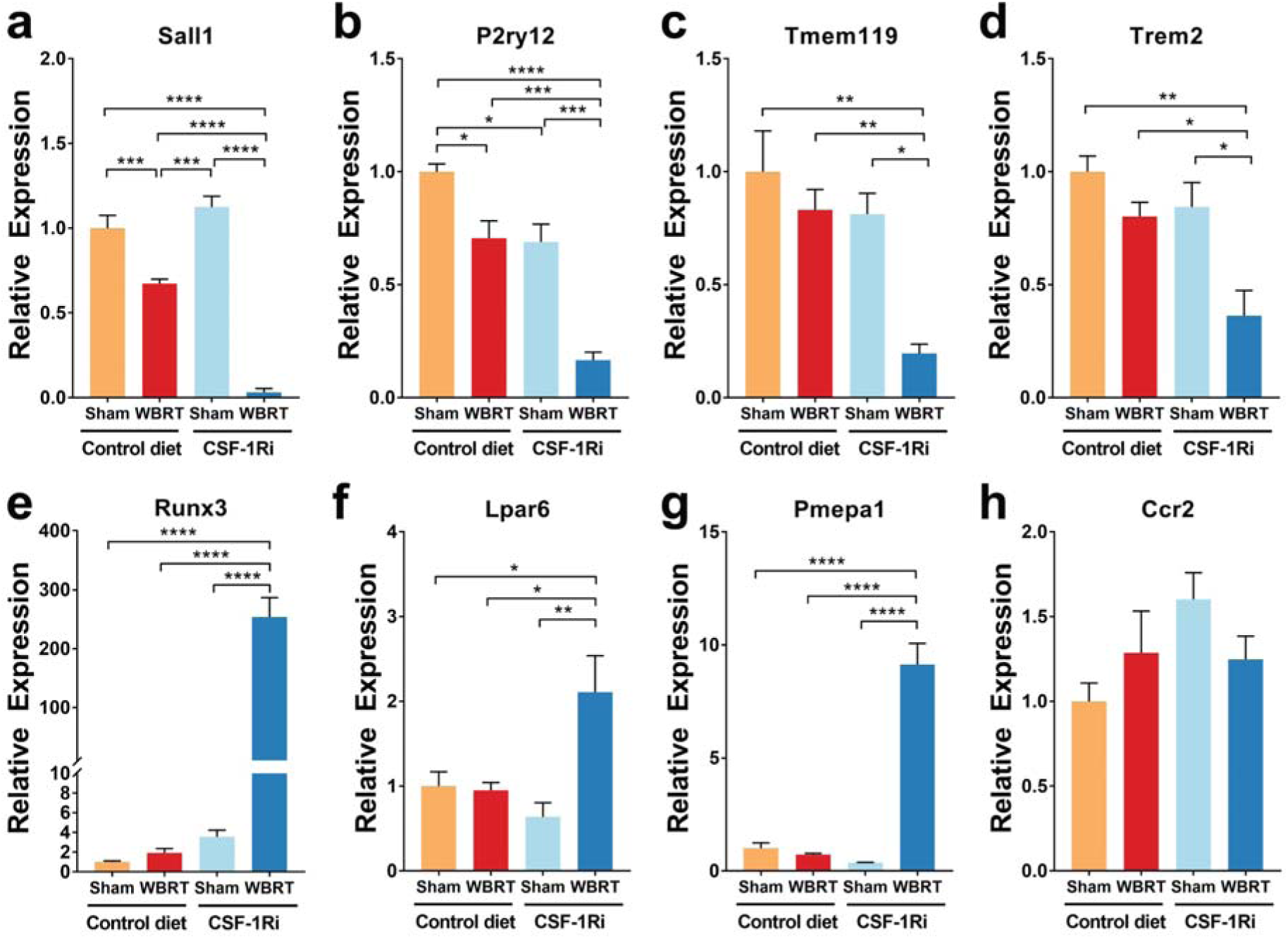
qPCR validation of microglia- and monocyte-specific genes. Selected genes that are known to highly express in microglia or monocytes were used to validate RNAseq results. **a – d** microglia signature genes *Sall1*, *P2ry12*, *Tmem119* and *Trem2* have lower expression levels in monocyte derived BEMs (CSF-1Ri + WBRT) compared to naïve microglia (control diet sham), irradiated microglia (control diet + WBRT) and repopulated microglia (CSF-1Ri sham). **e – f** monocyte signature genes *Runx3*, *Lpar6* and *Pmepa1* have higher expression levels in BEMs compared to other groups. Statistical analyses were performed using two-way ANOVA with Tukey’s multiple comparisons test. *p<0.05, **p<0.01, ***p<0.001, ****p<0.0001. N = 4 – 6. The qPCR experiments were performed in duplicates with similar results. Figures shown here are representative results from one experiment.

**Supplementary Figure 3:**
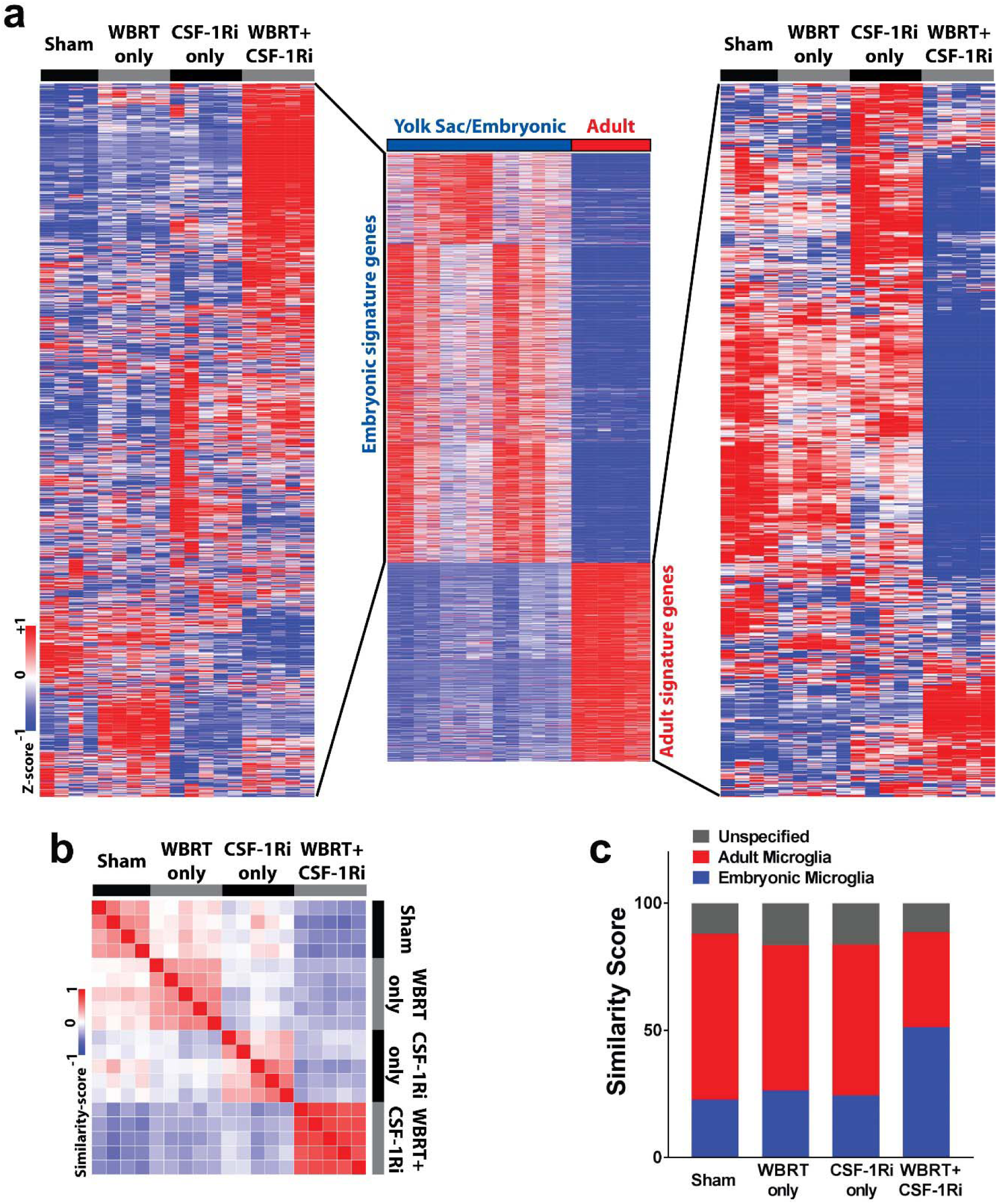
Monocyte-derived BEMs after WBRT have embryonic microglia signatures. **a** hierarchically clustered heatmaps to compare embryonic and adult microglia signatures across samples. Embryonic and adult signature genes were defined based on published dataset by Matchonitch and Winter et al. (Gene list and expression data in Supplementary Table 3). **b** Similarity matrix comparisons using defined embryonic and adult signature genes. **c** bar graph showing similarity scores to compare relative numbers of genes (shown as percentage of the defined list) that express in the same trends as embryonic or adult microglia in the Matchovitch and Winter dataset.

**Supplementary Figure 4:**
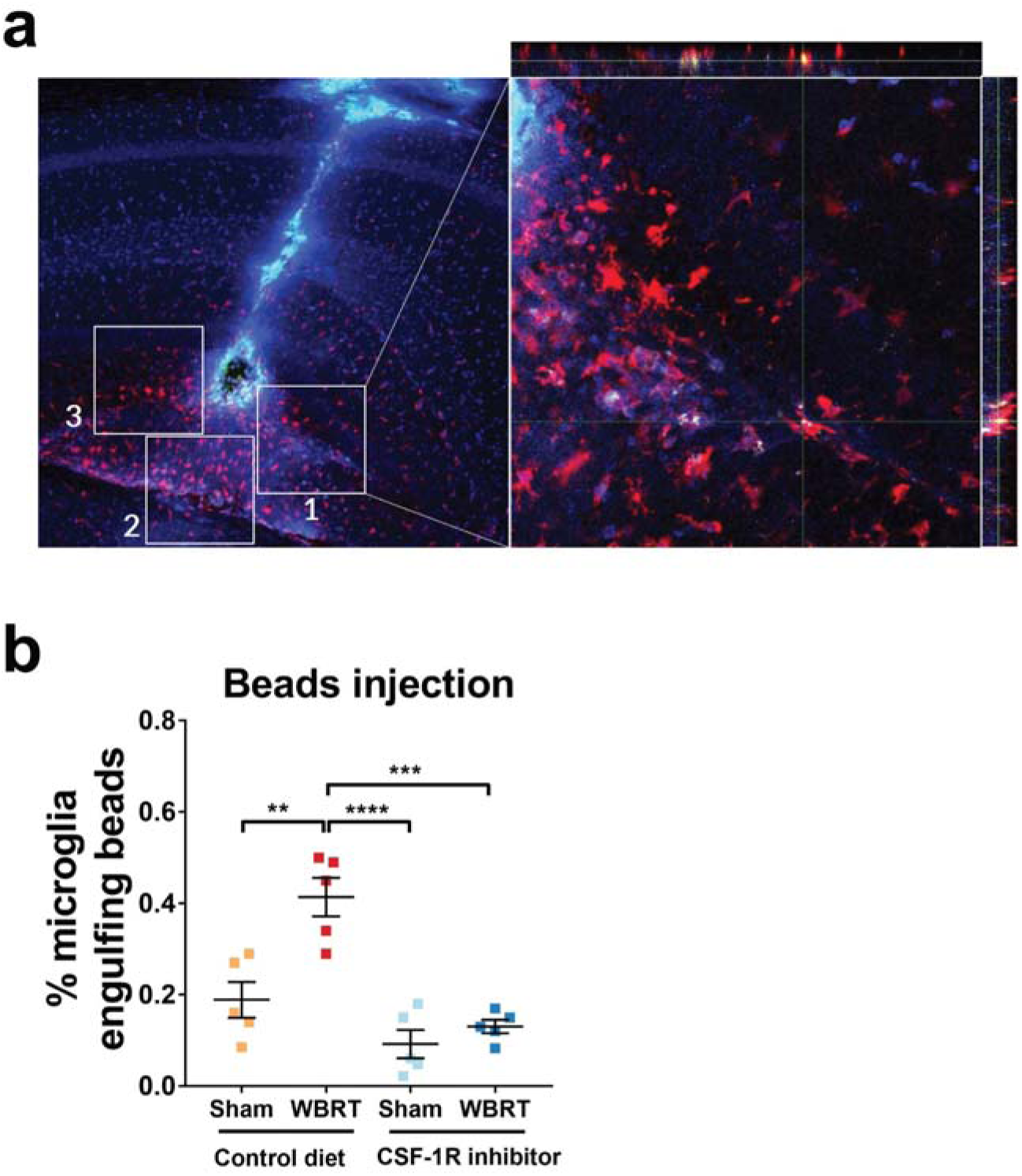
Representative images of count window from phagocytosis assay by IF and result of in vivo beads phagocytosis assay by FACS. A, representative images showing injection track of pre-stained synaptosomes and count windows. b, dot plot results of in vivo phagocytosis assay by FACS using fluorescent labeled beads. Statistical analyses were performed using two-way ANOVA with Tukey’s multiple comparisons test. **p<0.01, ***p<0.001, ****p<0.0001. N = 5.

**Supplementary Figure 5:**
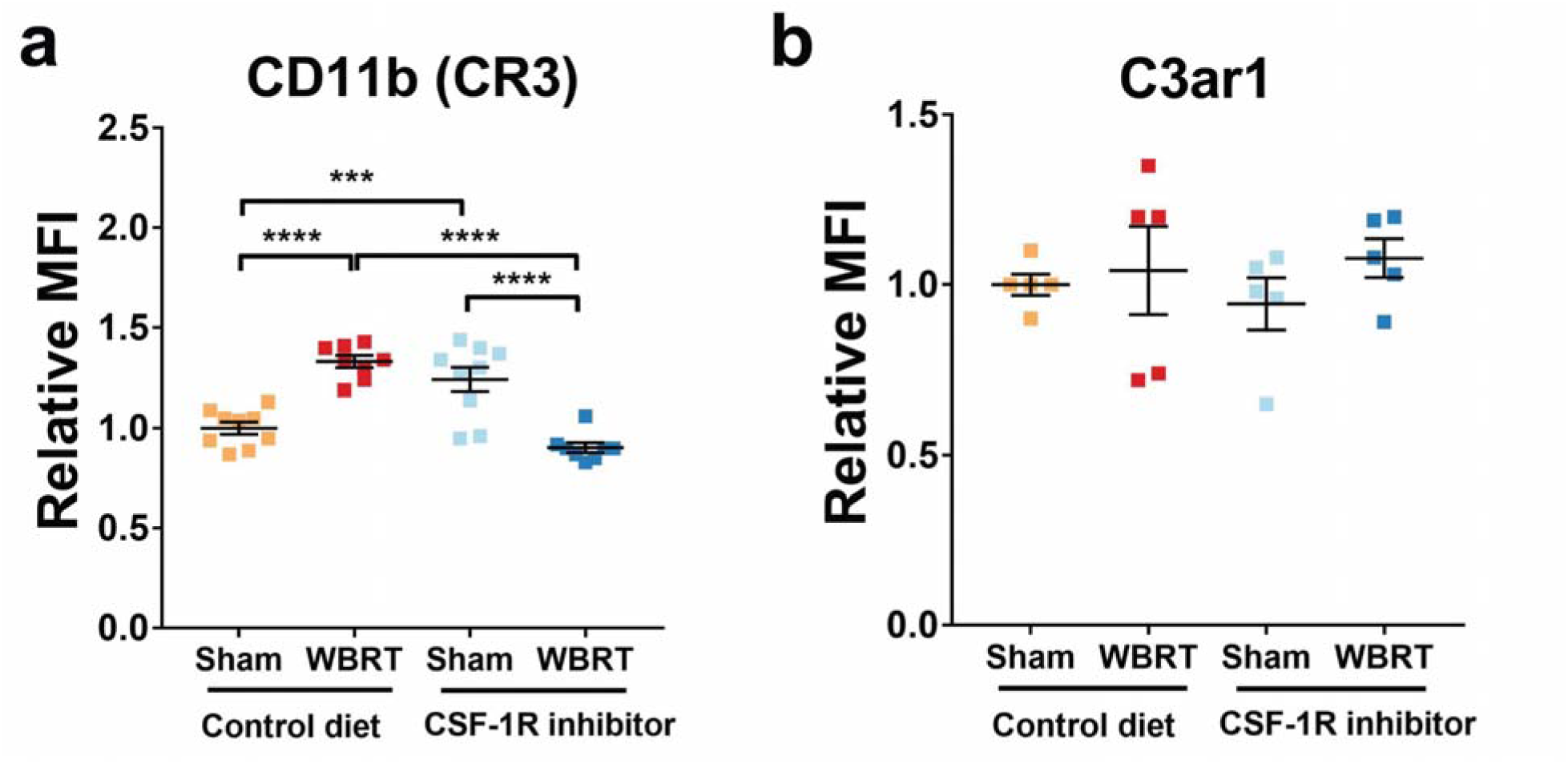
Complement receptors CR3 and C3ar1 levels in microglia and BEMs. a, dot plot of relative MFI of complement receptor CR3 (CD11b). b, dot plot of relative MFI of complement receptor C3ar1. Statistical analyses were performed using two-way ANOVA with Tukey’s multiple comparisons test. ***p<0.001, ****p<0.0001. N = 8 – 9 (a), N = 5 (b).

**Supplementary Figure 6:**
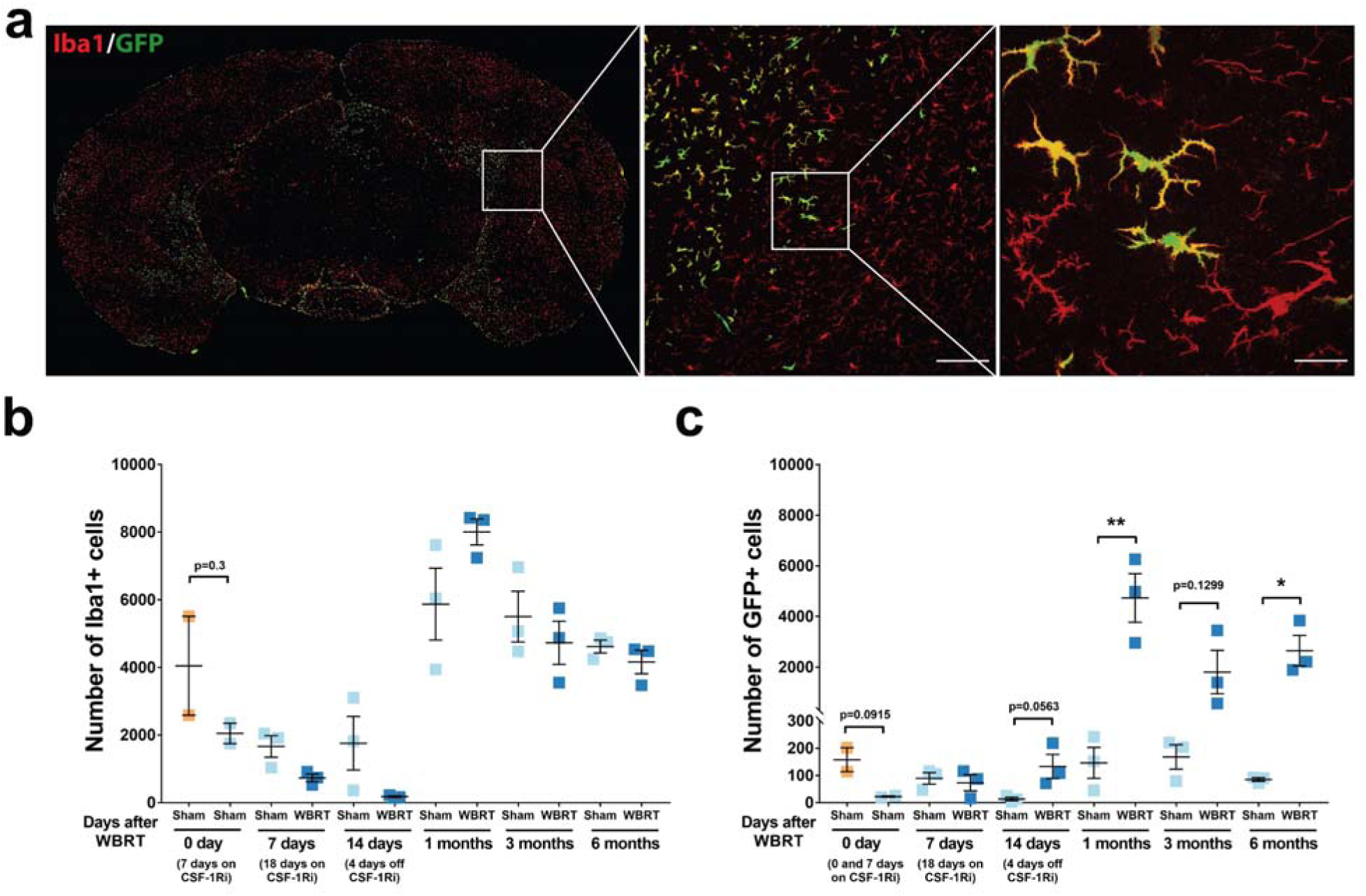
Quantification of microglia and BEMs in brains from bone marrow chimeras. **a** representative images of coronal section of whole brains from bone marrow chimeras. Scale bars = 100 μm (middle) and 20 μm (right). **b** dot plot of quantification results of Iba1 positive cells, each dot represents number of cells stained positive for Iba1 from a coronal whole brain section of an individual mouse. **c** dot plot of quantification results of GFP positive cells, each dot represents number of GFP positive cells from a coronal whole brain section of an individual mouse. Statistical analyses were performed using two-way ANOVA with Tukey’s multiple comparisons test. *p<0.05, **p<0.01. N = 2 (0 day sham on control diet) 3 (all other groups).

**Supplementary Figure 7:**
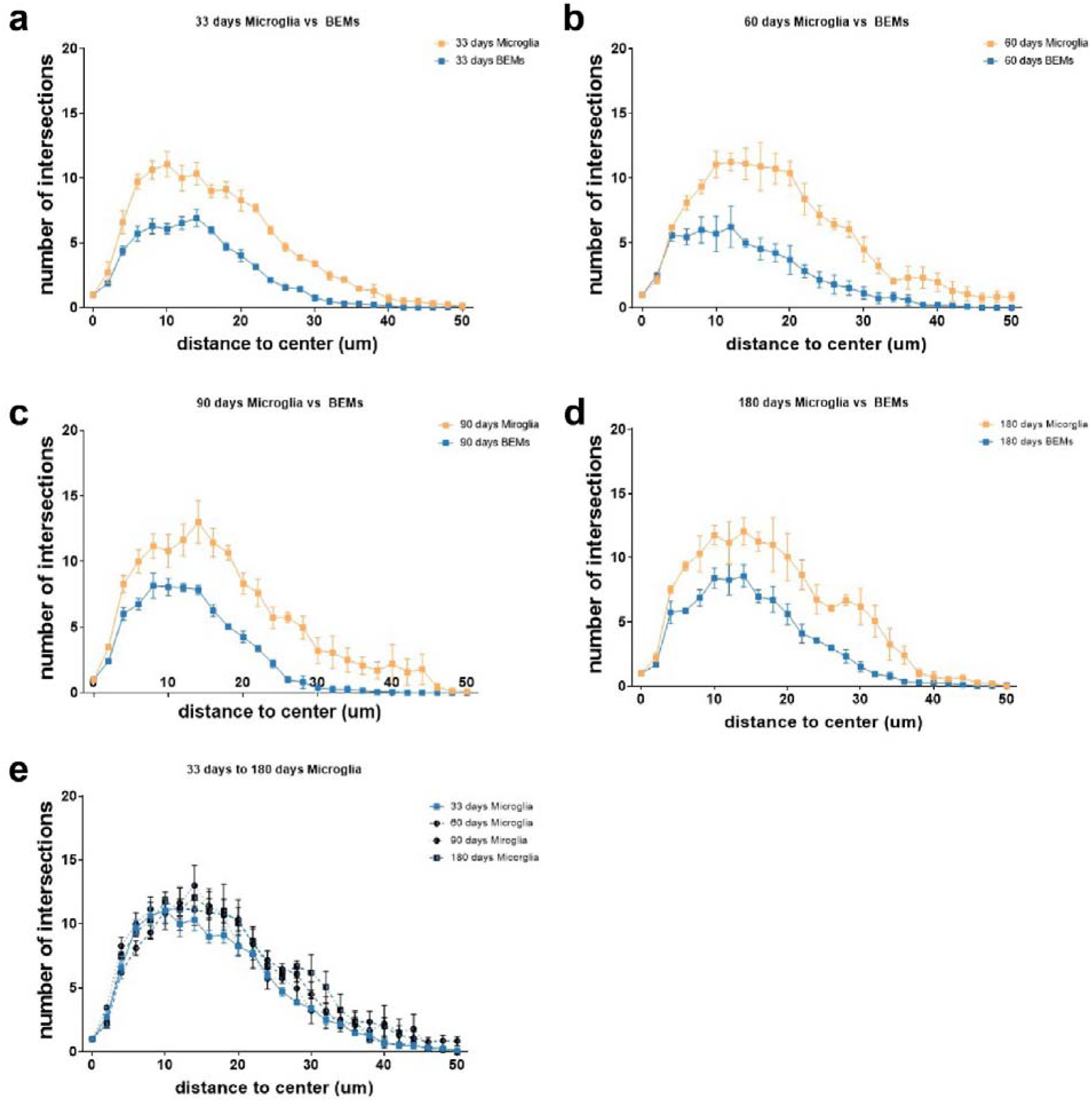
Sholl analysis results of microglia vs BEMs over time. a- d, comparison of Sholl analyses results between microglia and BEMs at 33, 60, 90 and 180 days after WBRT. e, Sholl analyses results of microglia at 33, 60, 90 and 180 days after WBRT. Statistics were performed using unpaired t-test at each distance point. n = 5 – 6.

